# Predicting Response to Platin Chemotherapy Agents with Biochemically-inspired Machine Learning

**DOI:** 10.1101/231712

**Authors:** Eliseos J. Mucaki, Jonathan Z.L. Zhao, Dan Lizotte, Peter K. Rogan

**Affiliations:** Department of Biochemistry, Schulich School of Medicine and Dentistry, Western University, London, Canada, N6A 2C1; Department of Computer Science, Faculty of Science, Western University, London, Canada, N6A 2C1; Department of Epidemiology & Biostatistics, Faculty of Science, Western University, London, Canada, N6A 2C1; Cytognomix Inc. London, Canada N5X 3X5; Department of Oncology, Schulich School of Medicine and Dentistry, Western University, London, Canada, N6A 2C1

**Keywords:** Chemotherapy response, support vector machines, gene signatures, cancer, cisplatin, oxaliplatin, carboplatin, machine learning, bladder cancer, breast cancer, ovarian cancer

## Abstract

Selection of effective genes that accurately predict chemotherapy response could improve cancer outcomes. We compare optimized gene signatures for cisplatin, carboplatin, and oxaliplatin response in the same cell lines, and respectively validate each with cancer patient data. Supervised support vector machine learning was used to derive gene sets whose expression was related to cell line GI_50_ values by backwards feature selection with cross-validation. Specific genes and functional pathways distinguishing sensitive from resistant cell lines are identified by contrasting signatures obtained at extreme vs. median GI_50_ thresholds. Ensembles of gene signatures at different thresholds are combined to reduce dependence on specific GI_50_ values for predicting drug response. The most accurate models for each platin are: cisplatin: *BARD1*, *BCL2*, *BCL2L1*, *CDKN2C*, *FAAP24*, *FEN1*, *MAP3K1*, *MAPK13*, *MAPK3*, *NFKB1*, *NFKB2*, *SLC22A5*, *SLC31A2*, *TLR4*, *TWIST1*; carboplatin: *AKT1*, *EIF3K*, *ERCC1*, *GNGT1*, *GSR*, *MTHFR*, *NEDD4L*, *NLRP1*, *NRAS*, *RAF1*, *SGK1*, *TIGD1*, *TP53*, *VEGFB*, *VEGFC;* oxaliplatin: *BRAF*, *FCGR2A*, *IGF1*, *MSH2*, *NAGK*, *NFE2L2*, *NQO1*, *PANK3*, *SLC47A1*, *SLCO1B1*, *UGT1A1*. TCGA bladder, ovarian and colorectal cancer patients were used to test cisplatin, carboplatin and oxaliplatin signatures (respectively), resulting in 71.0%, 60.2% and 54.5% accuracy in predicting disease recurrence and 59%, 61% and 72% accuracy in predicting remission. One cisplatin signature predicted 100% of recurrence in non-smoking bladder cancer patients (57% disease-free; N=19), and 79% recurrence in smokers (62% disease-free; N=35). This approach should be adaptable to other studies of chemotherapy response, independent of drug or cancer types.

## INTRODUCTION

Chemotherapy regimens are selected based on overall outcomes for specific types and subtypes of cancer pathology, progression to metastasis, other high-risk indications, and prognosis^1,2^, and variability in tumor resistance has led to tiered sequential strategies for selection of agents based on their overall efficacy^3^. We and others have developed machine learning (ML)-based gene signatures aimed at predicting response to specific chemotherapeutic agents and minimizing chemoresistance based on inhibition of growth or drug targets (GI_50_ or IC_50_)^4–6^. In this study, we present integrated ML models of platin drug responses (cis-, carbo- and oxaliplatin). Previous studies have reviewed the genes^7^, gene products^8^ and specific individual pathways that are activated and repressed by drugs^9^, but lack comprehensive models of the global cellular response to drugs. We use integrated ML-based signatures based on expression of multiple genes to predict key responses to each of these platin agents, for the first time, at different resistance levels.

Cisplatin, carboplatin and oxaliplatin are each widely prescribed compounds for their antineoplastic effects. While each contains platinum to form adducts with tumour DNA, their effectiveness differs for specific types of cancers, such as bladder (cisplatin), ovarian (cisplatin and carboplatin) and colorectal cancer (oxaliplatin). Carboplatin differs in structure from cisplatin, exchanging the latter’s dichloride ligands with a CBDCA (cyclobutane dicarboxylic acid) group, while oxaliplatin is paired with both a DACH (diaminocyclohexane) ligand and a bidentate oxalate group. These chelating ligands have greater stability and solubility to aqueous solutions, which lead to differences in drug toxicity compared to cisplatin^10^. Oxaliplatin can be up to two times as cytotoxic as cisplatin, but it forms fewer DNA adducts^11^. The large hydrophobic DACH ligand which overlaps the major groove is thought to prevent binding of certain DNA repair enzymes such as the POL polymerases, and may contribute to the low cross-resistance between oxaliplatin and cisplatin and carboplatin^10^. While all three drugs can enter the cell via copper transporters, organic cation transporters are oxaliplatin-specific and likely play a role in its efficacy in colorectal cancer (CRC) cells where these transporters are commonly overexpressed^7^. Oxaliplatin specifically plays a role in interfering with both DNA and RNA synthesis, unlike cisplatin which only infers with DNA^12^. It is these intrinsic properties between the platinum drugs which lead to differences in their activity and resistance profiles, despite their similar mode of action.

We derived gene signatures to predict drug response at different sensitivity and resistance levels for each of these agents. We and others have used supervised learning algorithms, including random forest models^13^; support vector machine (SVM) models^6^; neural networks^14^; and linear regression models^5^ to make these predictions. Pathway and network analysis of gene expression have been used to indicate hundreds of genes potentially up- and down-regulated upon cisplatin treatment^15^. Cisplatin-specific gene signatures have been developed with integrative approaches such as elastic net regression using inferred pathway activity of bladder cancer cell line data^16^. These methods have implicated genes that have not been described previously. Supervised ML with biochemically-relevant genes has also been useful for predicting drug response^6^. A concern with each of these ML approaches is that an insufficient number of samples coupled to a large number of features, i.e. gene expression changes, in each sample can result in overfitting of the model affecting its generalizability with other sources of data^17^. We therefore reduce the number of dimensions by selecting genes biologically relevant to the drugs under observation^6,17^. Additional selection criteria are necessary when the number of genes implicated in peer-reviewed reports is still prohibitively large compared to sample size.

Biochemically-inspired gene signatures have shown good performance in predicting treatment response. A paclitaxel ML signature based on tumor gene expression had a higher success predicting the pathological complete response rate (pCR ^18^) for sensitive patients (84% of patients with no / minimal residual disease) than models based on differential gene expression (GE) analysis^6^. For gemcitabine, a signature derived from both expression and copy number (CN) data from breast cancer cell lines was derived, and subsequently applied to analysis of nucleic acids from patient archival material. Multiple other outcome measures used to validate gene signatures include prognosis^5^, Miller-Payne response^19^, and disease recurrence. Binary SVM classifiers based on discrete time thresholds have been used to classify continuous outcome measures such as prognosis and recurrence. By contrast, pCR is simpler to interpret with binary SVM models. Nevertheless, differences in clinical recurrence have been noted between patients demonstrated with pCR and those who do not exhibit disease pathology^18^. This source of variability in defining patient response can confound transferability of SVM models between different datasets.

We apply biochemically-inspired ML to predict and compare the cellular and patient responses to cisplatin, carboplatin and oxaliplatin. We train models for classification of platin resistance with cancer cell line data and validate with patient GE and outcome data. Our previous gene signatures were based on median GI_50_ for each drug^6^. This has been a necessary compromise, however in this study we consider signatures that differ at the highest vs. the lowest levels of drug resistance. A series of gene signatures are derived by shifting the GI_50_ thresholds that distinguish sensitivity from resistance. The frequency of genes selected at median vs. extreme thresholds highlights pathways that define these responses among different patient subsets.

## RESULTS

### Selection of Platin Drug Related Genes

We documented genes in the peer-reviewed literature associated with drug effectiveness or response (Supplemental References). For cisplatin, carboplatin and oxaliplatin, this implicated 178, 90, and 288 genes, respectively (Suppl. Table S1). Multiple factor analysis (MFA) was used to determine which genes were correlated to GI_50_ in breast cancer cell lines through either GE and/or CN^13^, significantly reducing the sizes of the gene sets for cisplatin (N=39), carboplatin (N=28), and oxaliplatin (N=55). Genes with significant relationships to GI_50_ and direction of correlation (positive or inverse) are indicated in Figure 1. The diverse functions of these genes included apoptosis, DNA repair, transcription, cell growth, metabolism, immune system, signal transduction and membrane transport. Analysis of IC_50_ and gene expression levels for cisplatin-treated bladder cancer cell lines confirmed these relationships evident from GI_50_ values of different breast cancer lines. IC_50_ values were related to GE for *CFLAR*, *FEN1*, *MAPK3*, *MSH2*, *NFKB1*, *PNKP*, *PRKAA2*, and *PRKCA*^20^. Similarly, separate bladder cell line IC_50_ values from the Genomics of Drug Sensitivity in Cancer project (http://www.cancerrxgene.org; N=17) were correlated with GE for *CFLAR*, *FEN1*, and *NFKB1*, in addition to *ATP7B*, *BARD1*, *MAP3K1*, *NFKB2*, *SLC31A2* and *SNAI1*.

**Figure 1.**
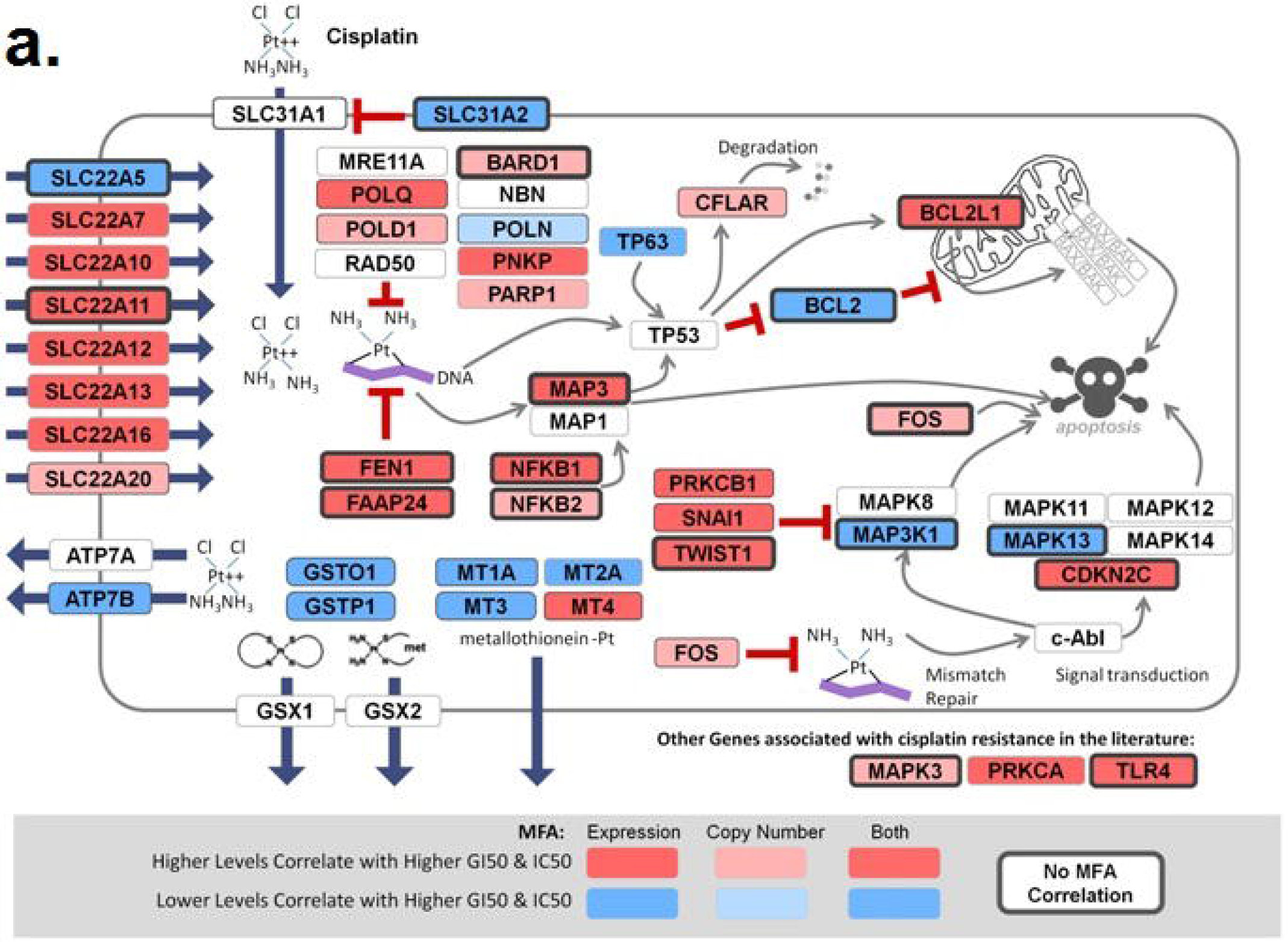
Schematic of platinum drug sensitivity and resistance genes which showed MFA correlation for GI_50_ of A) cisplatin, B) carboplatin, and C) oxaliplatin. The genes used to derive the SVM are shown in context of their effect in the cell and role in cisplatin mechanisms of action. GE and CN correlation with inhibitory drug concentration by MFA of breast (GI_50_) and bladder (IC_50_) cancer cell line data.

We performed MFA on the GI_50_ values for cisplatin, carboplatin and oxaliplatin, without consideration of either GE or CN. Responses to cis- and carboplatin were directly correlated (a 6.2º separation between vectors), but neither was related to the oxaliplatin response (Figure 2). Previous studies have shown that cisplatin-resistant cell lines are generally sensitive to oxaliplatin^21–23^.

**Figure 2.**
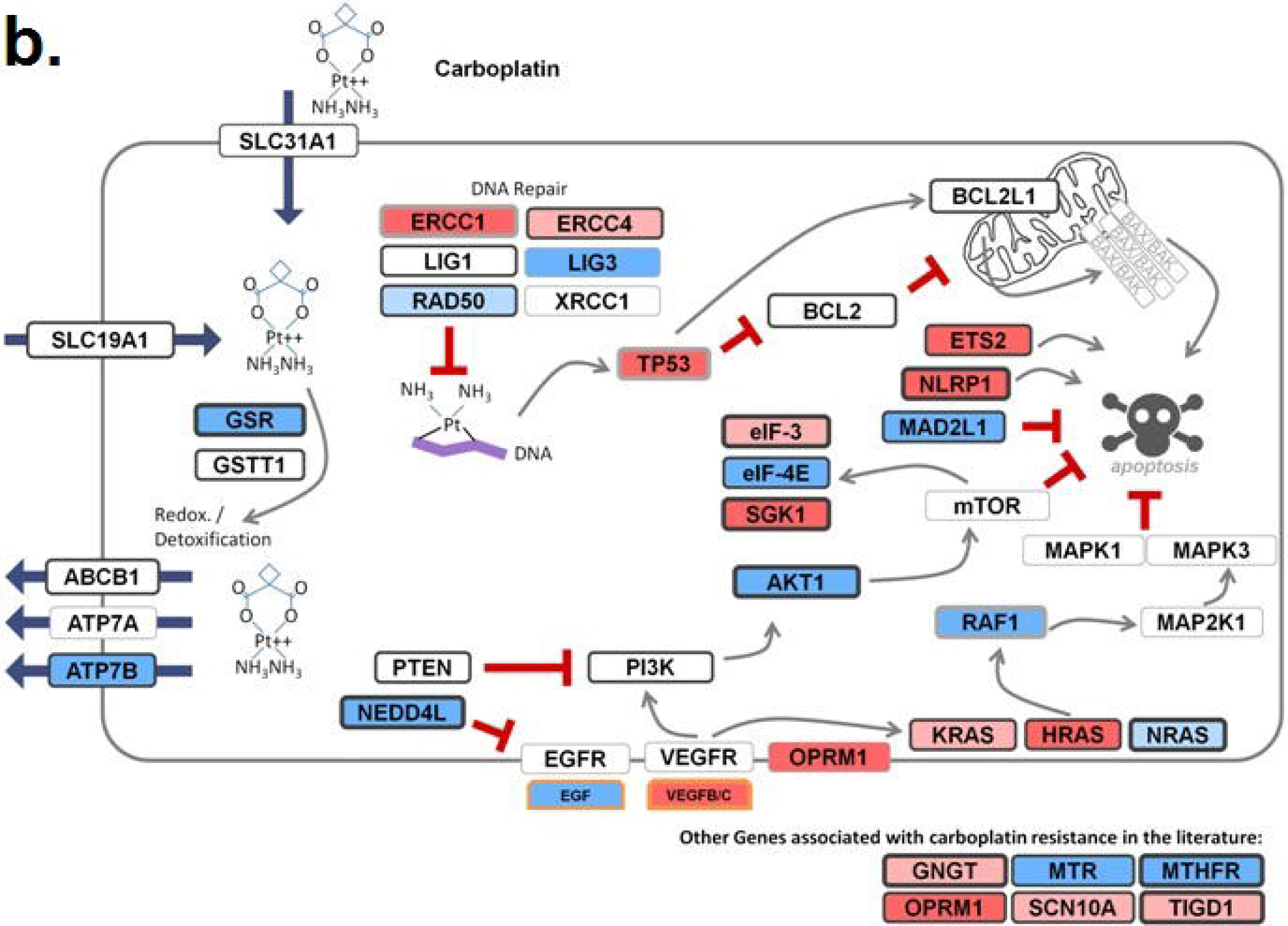
GI_50_ values for cell lines treated with the three platin drugs were plotted in order of ascending oxaliplatin GI_50_. For most cell lines, there is a visible trend between the GI_50_ for cisplatin and carboplatin, reflecting the correlation between the two drugs seen by MFA. Despite this correlation, carboplatin shows a much smaller variance (0.22) compared to cisplatin (0.37; oxaliplatin variance is 0.34).

SVM-based signatures were initially derived for each platin drug from breast cancer cell line GE data. A 13-gene signature for cisplatin at the median GI_50_ threshold (5.2% misclassification rate) consisted of *BARD1, BCL2L1, FAAP24, CFLAR, MAP3K1, MAPK3, NFKB1, POLQ, PRKAA2, SLC22A5, SLC31A2, TLR4,* and *TWIST1*. A similarly derived carboplatin signature included *AKT1*, *ATP7B*, *EGF*, *EIF3I*, *ERCC1*, *GNGT1*, *HRAS*, *MTR*, *NRAS*, *OPRM1*, *RAD50*, *RAF1*, *SCN10A*, *SGK1*, *TIGD1*, *TP53*, and *VEGFB* (10.4% misclassification). For oxaliplatin, the final SVM model consisted of *AGXT*, *APOBEC2*, *BRAF*, *CLCN6*, *FCGR2A*, *IGF1*, *MPO*, *MSH2*, *NAGK*, *NAT2*, *NFE2L2*, *NOTCH1*, *PANK3*, *PRSS1*, and *UGT1A1* (2.1% misclassification). A cisplatin SVM generated from 17 bladder cancer cell lines in cancerRxgene resulted in 2 equally accurate signatures (with 11.8% misclassification) consisting of either *PNKP* and *PRKCA* or *ATP7B*, *CFLAR*, *FEN1*, *MAPK3*, *NFKB1* and *SLC22A11*. These models were not useful for predicting patient outcomes due to the limited size of the training set.

### GI_50_-Threshold Independent Modeling

In our previous studies, we set median GI_50_ value as the threshold to distinguished drug resistance and sensitivity^5,6^. An important question is whether the genes contributing to drug response are consistent among different cell lines, each with their own unique GI_50_ values. Different ML models were obtained by shifting the GI_50_ threshold, which changed the labels of resistant vs. sensitive cell lines. After feature selection, the compositions of the corresponding gene signatures for each threshold were compared. Finally, ensemble averaging of all of these optimized Gaussian SVM models derived for different GI_50_ thresholds was used to create a threshold-independent ML-based signature.

Kinase (*MAPK3*, *MAP3K1*) genes and apoptotic family members (*BCL2*, *BCL2L1*) were most the common in the cisplatin signatures at different GI_50_ thresholds, with consistent representation of error-prone and base-excision DNA repair genes as well (Figure 3A; Supplementary Table S2A). The kinases are more concentrated in signatures with lower drug sensitivity thresholds, whereas *BCL2* and *BCL2L1* are more ubiquitous at all levels. The error prone polymerases, *POLD1* and *POLQ,* are more frequent in models with lower sensitivity thresholds, while the flap endonuclease *FEN1* tends to be present at high levels of resistance. Thresholded models for carboplatin-related genes commonly contained the apoptotic family member *AKT1*, transcription regulation genes *ETS2* and *TP53*, as well as cell growth factors *VEGFB* and *VEGFC*, although the latter was less common at lower sensitivity thresholds (Figure 3B). Common oxaliplatin-related genes included transporters *SLCO1B1* and *GRTP1* (but not *SLC47A1*), transcription genes *NFE2L2*, *PARP15* and *CLCN6*, as well as multiple metabolism-related genes (Figure 3C).

**Figure 3.**
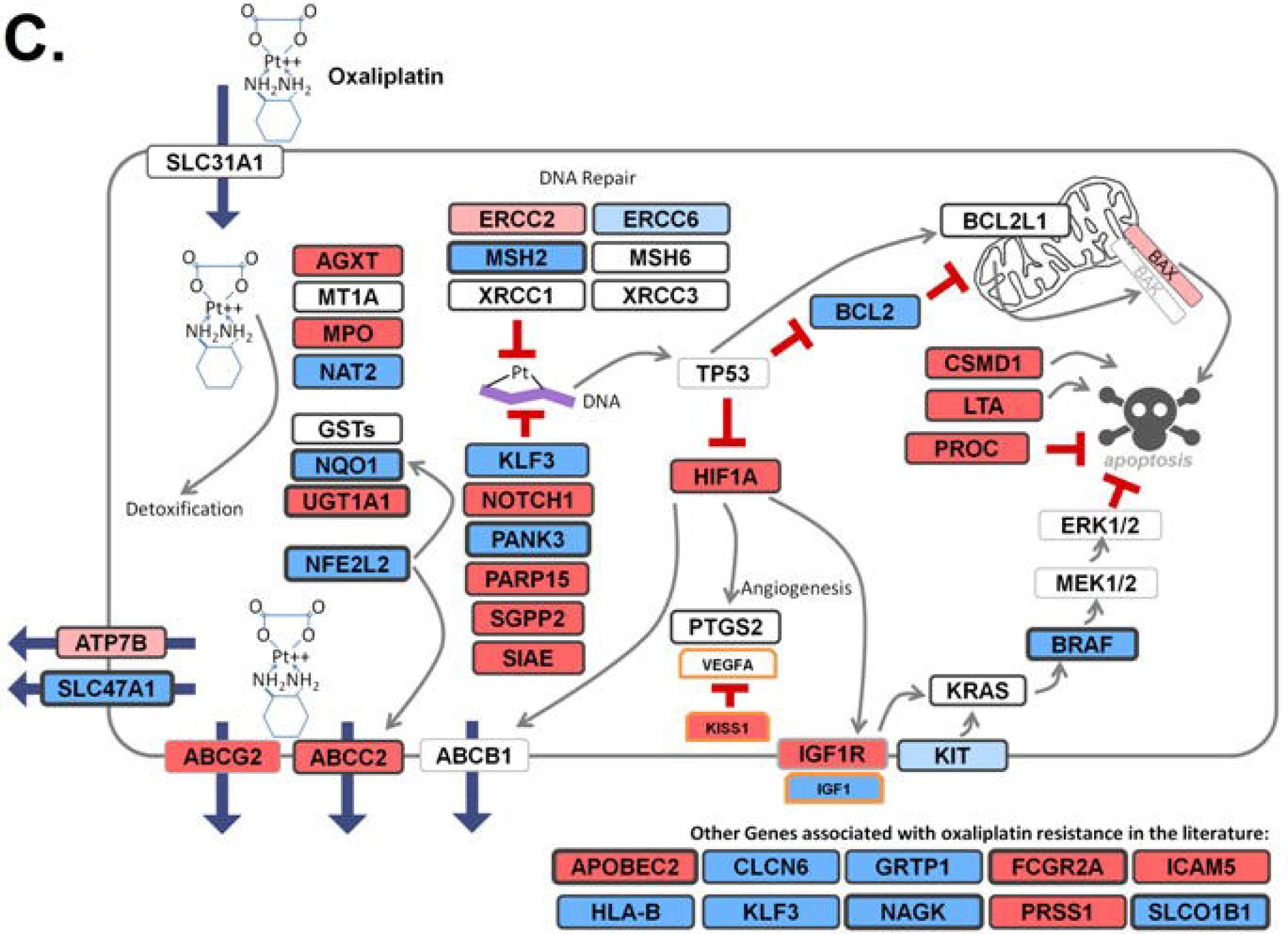
The variation in gene composition of misclassification-based SVMs at different GI_50_ thresholds for A) cisplatin, B) carboplatin, and C) oxaliplatin. GI_50_ intervals are indicated on the left, with the number of cell lines with GI_50_ values within said intervals in brackets. Each box represents the density of genes appearing in optimized Gaussian SVM models in those functional categories, with darker grey indicating frequent genes in indicated GI_50_ threshold intervals, while lighter grey indicates less commonly selected genes. The number of thresholded models used to derive the density plot within each interval is equal (or greater, in the case of multiple equally performing models) to the number of cell lines within that GI_50_ interval.

SVM models were also derived using the cisplatin and/or carboplatin-treated TCGA (The Cancer Genome Atlas) bladder urothelial carcinoma patients, using post-treatment time to relapse as a surrogate criterion for different GI_50_ resistance thresholds (as performed in Mucaki *et al.* [2017]^24^; Supplementary Table S3). Similar trends to cell line SVMs are apparent: *POLQ* is frequently included in models with recurrence threshold of longer duration, while *FEN1* is a marker of resistance, when time to relapse is shorter. However *BCL2*, which is present in a majority of breast cancer cell line SVMs, is present in only one model derived from TCGA data. Similarly, *MSH2* was rarely selected using cell lines, yet appears in nearly all patient derived SVMs with > 1 year recurrence.

GI_50_ thresholded ML models were also derived using the log-loss function, which penalizes false classifications, whose value ranges from zero (or completely accurate), to 1 (or completely inaccurate; Supplementary Table S4). The overall distribution of genes across GI_50_ thresholds has many distinct similarities with the models derived by misclassification. For both sets of cisplatin models, *BCL2*, *BCL2L1* and *FEN1* are common in low-to-moderate GI_50_ thresholds, while *NFKB1* is enriched at high thresholds (Figure 3A; Suppl. Figure 1A). For carboplatin, *AKT1*, *VEGFB* and *VEGFC* are similarly distributed across GI_50_ thresholds with both methods, although *VEGFB* is less dense with log-loss models at low GI_50_ values (Figure 3B; Suppl. Figure 1B). In both sets of models for oxaliplatin, *SIAE* and *SLC47A1* show a high density across all GI_50_ thresholds, while *ABCG2* shows low density across each (<50% inclusion; Figure 3C and Suppl. Figure 1C). There are some distinct differences. *EGF* and *ERCC1* were selected at a greater frequency at a moderate carboplatin GI_50_ with log-loss rather than misclassification. Similarly, the following oxaliplatin genes were selected considerably more often when using log-loss: *APOBEC2*, *HLA-B*, *LTA*, and *MPO*. Therefore, while the misclassification and log-loss based models are not interchangeable, the models are overall quite similar.

Log-loss models were initially constructed either by (a) a modified version of the misclassification-based method, or (b) using the BFS software described in Zhao *et al.* (2018)^25^. Multiple signatures with low log-loss values can have different compositions, consistent with the possibility that there may be various diverse gene combinations that can give rise to signatures with satisfactory performance. However, these signatures often contain a larger number of gene features than the misclassification based signatures, and raised concerns that they might be more prone to overfitting. The log-loss minimized models generated by both methods had comparable compositions. The median GI_50_ thresholded cisplatin model generated by the log-loss modified software [*ATP7B*, *BCL2L1*, *CDKN2C*, *CFLAR*, *ERCC2*, *ERCC6*, *FAAP24*, *FOS*, *GSTO1*, *GSTP1*, *MAP3K1*, *MAPK13*, *MAPK3*, *MSH2*, *MT2A*, *PNKP*, *POLD1*, *POLQ*, *PRKAA2*, *PRKCA*, *PRKCB*, *SLC22A5*, *SLC31A2*, *SNAI1*, *TLR4*, *TP63*] shares 15/19 genes with the signature generated by the BFS software^25^ [*ATP7B*, *BARD1*, *BCL2*, *BCL2L1*, *ERCC2*, *FAAP24*, *FEN1*, *FOS*, *MAP3K1*, *MAPK13*, *MAPK3*, *MSH2*, *MT2A*, *NFKB1*, *PNKP*, *POLQ*, *PRKCB*, *SLC22A5*, *SNAI1*]).

### Traditional Model Validation against Cancer Patient Data

GI_50_-thresholded models for each platin drug, generated with the breast cancer cell line data, produced 70 cisplatin, 83 carboplatin, and 83 oxaliplatin SVM models, respectively. Each model was validated using available platin-treated patient datasets^26–30^. The chemotherapy response metadata differed between studies. Als *et al.*^29^ reported survival post-treatment, whereas Tsuji *et al.*^30^ categorized patients as responders and non-responders. TCGA provided two different measures which were used to assess predictive accuracy in our models – chemotherapy response and disease-free survival. Accuracy is similar using either measure (Supplementary Table S5A); however recurrence and disease-free survival was used as the primary measure of response as it was more often recorded in the TCGA data sets tested. Patients from Als *et al.* with a ≥ 5 year survival post-treatment were labeled as sensitive to treatment. The differences between these metadata may, in part, account for the differences in the prediction accuracy of the thresholded SVM models.

At higher resistance thresholds for any platin drug (low GI_50_), where more cell lines are labeled sensitive, the positive class (disease-free survival) is correctly classified, while the negative class (recurrence) is highly misclassified (Suppl. Figures 2 and 3). The reverse is true for models built using lower resistance thresholds (high GI_50_). We therefore state SVMs generated at these extreme thresholds are not very useful at predicting patient data. When used to predict recurrence in the TCGA datasets, sensitivity and specificity appears to be maximized in models where the GI_50_ threshold for resistance was set near (but not necessarily at) the median (Suppl. Figure 2; Suppl. Tables S5A to 5C). While this pattern holds true for Tsuji *et al.*^30^, oxaliplatin models where GI_50_ thresholds were set above the median could better separate primary and metastatic CRC patients (best model predicting 92.6% metastatic and 60.7% primary cancers; Suppl. Table S5C). While less consistent, cisplatin models generated with thresholds above median GI_50_ performed better when evaluating the Als *et al.*^29^ patient dataset (Suppl. Figure 3).

Models were further evaluated for their accuracy in TCGA patients using various recurrence times post-treatment to classify resistant and sensitive patients (0.5 - 5 years; Supplemental Table S6A-C). The best performing cisplatin model (hereby identified as **Cis1**; Table 1) was able to accurately predict 71.0% of bladder cancer patients who recurred after 18 mo. (N=31; 58.5% accurate for disease-free patients [N=41]). Response of TCGA bladder patients treated with carboplatin (without cisplatin; N=19) were best predicted by **Cis12** two years post-treatment (80% accurate for responding patients [N=5]; 93% for recurrent patients [N=14]). The best performing carboplatin model (designated **Car1** [Table 1]) predicted recurrence of ovarian cancer after 4 years at an accuracy of 60.2% (N=302; 61.0% accurate for disease-free patients [N=108]). These models were also used to test TCGA bladder cancer patients treated carboplatin but not cisplatin (N=19), of which the best performing model (**Car73**) was 84% accurate for patients after 1 year of treatment (100% for responding patients [N=11]; 62.5% accuracy for recurrent [N=8]). Two additional carboplatin models are tied for overall accuracy (84%; **Car9** and **Car51**), but more successfully predict non-responsive patients (87.5%; 82% accuracy for responding patients). These three models share four genes: *AKT1, ETS2, GNGT1*, and *VEGFB.* For oxaliplatin, the best performing model (designated **Oxa1** [Table 1]) accurately predicted 71.6% of the disease-free TCGA CRC patients after one year (N=88; 54.5% accuracy predicting recurrence [N=11]). These models (based on gene expression measured by Affymetrix Gene Chip Human Exon 1.0 ST arrays), TCGA sample expression data, as well as SVMs based on bladder cell line data (based on expression measured by Affymetrix U133A microarray), were added to the online web-based SVM calculator (http://chemotherapy.cytognomix.com; introduced in Dorman *et al.* [2016]^6^) to predict platin response.

**Table 1:**
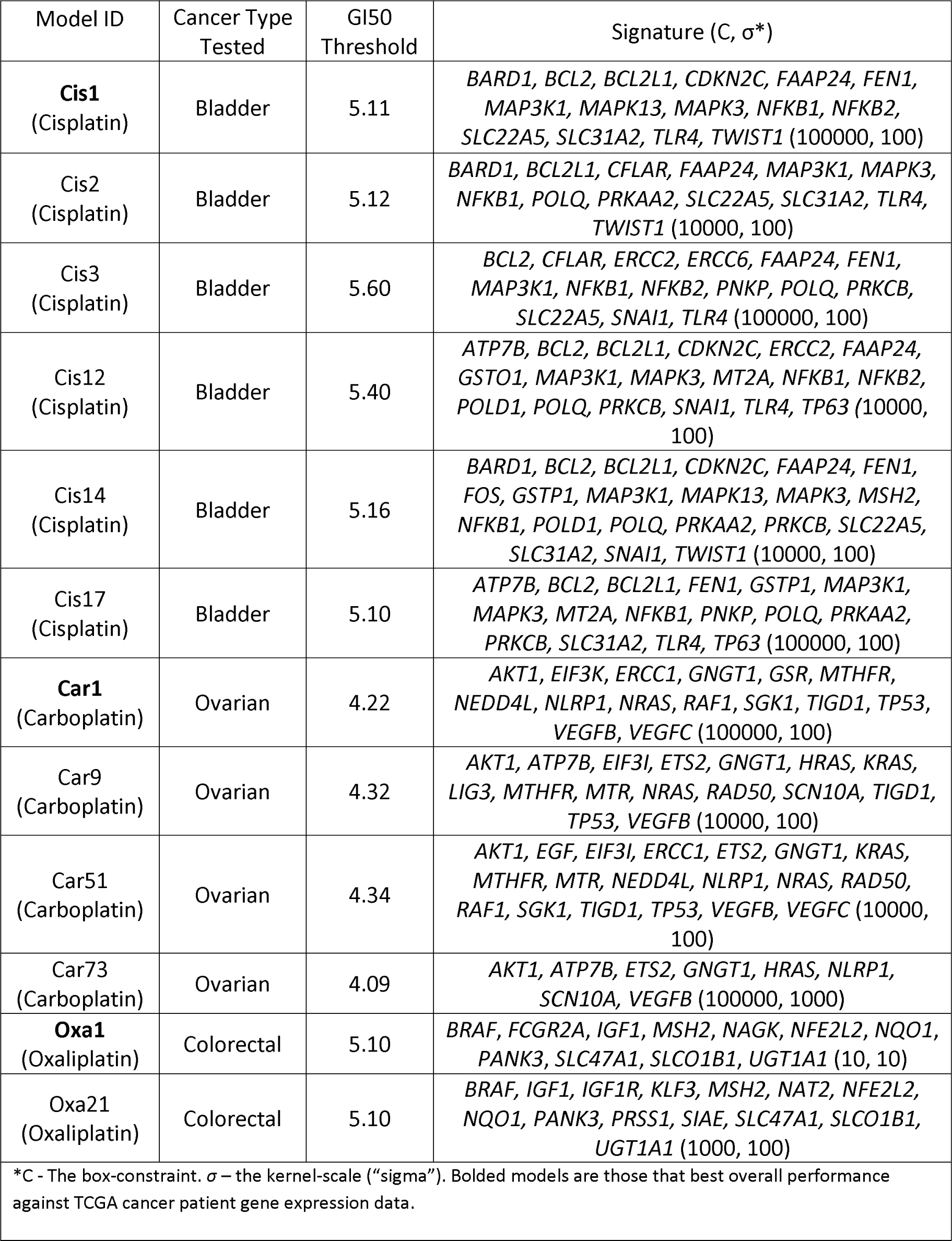
Models Which Best Predicted Response in TCGA Cancer Patients

To evaluate the consistency in the response prediction of TCGA bladder cancer patients treated with cisplatin, the distance from the hyperplane for all SVMs generated were plotted for each patient with a short recurrence time (<6 mo., N=10; Supplementary Figure 4). Despite showing similar levels of resistance to treatment, patterns differed between patients. While these patients would be expected to be indicated as highly cisplatin resistant (hyperplane distance < 0), two patients (TCGA-XF-A9SU and TCGA-FJ-A871) were predicted sensitive across nearly all SVM models. Similar variation was also seen in patients with either a long recurrence time (>4 years) or no recurrence at all after 6 years (Suppl. Figure 5).

Threshold independent models were generated for each individual platin drug at different GI_50_ thresholds through ensemble ML, which involves the averaging of hyperplane distances for each model to generate a composite score for each TCGA patient tested. Hyperplane distances across all 70 cisplatin models were similar, with a mean score of -0.22 and a standard deviation of 3.5 hyperplane units (hu) across the set of patient data. The ensemble model classified disease-free bladder cancer patients with 59% accuracy and those with recurrent disease with 47% accuracy. Limiting ensemble averaging to only cisplatin models generated at a moderate GI_50_ threshold (ranging from 5.10 to 5.50) did not significantly improve accuracy (44% for disease-free and 66% for recurrent patients; Suppl. Table S7A). For carboplatin, ensemble ML did not produce significantly better predictions than random, regardless of the GI_50_ threshold interval selected (Suppl. Table S7B) or the similar mean hyperplane distances (-0.11 +/- 3.9 hu). For oxaliplatin, the ensemble ML model (mean = -0.12 +/- 2.7 hu) was most accurate after 1 year (60% accuracy for disease-free and 73% for recurrent patients; Suppl. Table S7C). As in cisplatin, limiting this analysis to oxaliplatin SVM models with moderate GI_50_ thresholds did not significantly increase accuracy.

To determine the impact of individual genes on overall model accuracy, each gene within every SVM model was excluded, and model accuracy was reassessed (Supplementary Tables S2A; S2B and S2C contain cis-, carbo- and oxaliplatin models, respectively). Genes which consistently significantly increase misclassification (averaging > 16% increase) in moderate threshold SVMs (GI_50_ thresholds set from 5.1 to 5.5) include *ERCC2*, *POLD1*, *BARD1*, *BCL2*, *PRKCA* and *PRKCB*. *ERCC2* and *POLD1* perform critical functions in nucleotide and base excision repair, respectively. *PRKCA* and *PRKCB* are paralogs with significant roles in signal transduction. *BARD1* has been shown to reduce apoptotic *BCL2* in the mitochondria^31^, and has a key role in genomic stability through its association with *BRCA1*. Genes with a high variance in increased misclassification between different models include *NFKB1*, *NFKB2*, *TWIST1*, *TP63*, *PRKAA2*, and *MSH2*. The variance of these genes may be due to epistatic interactions with other biological components, including the other genes in the SVM. For example, *NFKB1* and *NFKB2* are jointly included in 7 SVMs generated at a moderate GI_50_ threshold. There is evidence of possible epistasis in that the removal of either of these genes, but not necessary both, will have a large impact in model misclassification rates (≥ 18.0% increase). The misclassification variance of *NFKB1* with *NFKB2*, is significantly lower than in SVM models lacking *NFKB2*.

To further evaluate the predictive capability of the misclassification-based gene signatures, k-fold cross-validation of the cisplatin, carboplatin and oxaliplatin models were performed on TCGA bladder, ovarian and colorectal cancer patient data, respectively. Patients were evenly distributed in 5 groups with an equal (or near-equal) ratio of disease-free and recurrent patients. The majority of the cisplatin models showed an overall accuracy > 50%. The cisplatin model which performed best under the k-fold analysis (6-resistance level; *BARD1*, *BCL2*, *BCL2L1*, *PRKAA2*, *PRKCA*, *PRKCB*, *TWIST1*) showed an overall accuracy of 71.2% (84.4% accurate for sensitive and 53.9% accurate for resistant patients). The accuracy of the carboplatin and oxaliplatin models did not exceed 60%. In general, traditional validation outperformed the k-fold validation results.

### Predicting cisplatin response in patients based on smoking history

Tobacco smoking is known as the highest risk factor for the development of bladder cancer^32^. We therefore subdivided the patients based on their smoking history and tested the thresholded models (Supplementary Tables S8 and S9). When testing patients who were lifelong non-smokers, the prediction accuracy of **Cis1** predicted all non-smoking patients who were recurrent after 18 months as cisplatin-resistant (N=5). Prediction accuracy for disease-free patients was 57.1% (N=14). Another model (**Cis18**; Suppl. Table S8) had performed equally as well for non-smokers, and these two models share 7 genes: *BCL2*, *BCL2L1*, *FAAP24*, *MAP3K1*, *MAPK13*, *MAPK3*, and *SLC31A2*. Threshold independent analysis predicted disease-free equally well, but recurrence was less accurate (66.7%). Note that non-smokers make up a small subset of the patients tested (N=19). Threshold-independent prediction of recurrence in patients with a smoking history was 46% accurate (N=13), while disease-free patients were correctly predicted at a rate of 58% (N=19). Recurrence in these patients was best predicted by a model built at the median GI_50_ threshold (**Cis2**). Accuracy improved for both disease-free (57.7% -> 61.9%) and recurrent patients (76.0% -> 78.6%) when excluding patients who quit smoking more than 15 years before diagnosis. Genes in this SVM which are not present in the two models which performed well for non-smokers include *CFLAR* and *PRKAA2*.

Tobacco smoking has a significant impact on cytosine methylation levels in the genome^33^. CpG island methylation has been associated with smoking pack years in a subset of the TCGA bladder urothelial carcinoma patients^26^. We suspected that the level of methylation measured in the SVMs which performed best for smoking and non-smoking patients might differ, and with possible concomitant effects on GE. When ranking each gene from **Cis1** by highest methylation and GE, 88 of 1080 patient: gene combinations showed the expected inverse correlation between methylation levels and GE (i.e. high methylation and low GE). Inverse correlation of methylation and GE was more common than direct correlation (i.e. high methylation and high GE; N=17). However, direct correlation was more common in patients with a recent smoking history (70.5%). This pattern was also observed for **Cis2**, which best predicted recurrence in smokers. In cases where methylation and GE are directly correlated, we propose that smoking may alter expression by other effects, e.g. mutagenic, rather solely than by epigenetic inactivation through methylation.

To determine which genes in these models led to discordant predictions of patient outcome, we conducted a bioinformatic analysis in which the expression of each signature gene was gradually altered until the misclassification was corrected. If the GE value required to cross this threshold exceeded ≥ 3-fold the highest/lowest expression of that gene, it was interpreted as a minor contributor to the prediction. Genes which could not correct a discordant prediction included *PRKAA2*, *NFKB1*, *NFKB2* and *TWIST1*. Significant genes which, when altered, corrected discordant predictions included *MAP3K1*, *MAPK3*, *SLC22A5* and *SLC31A2*. Altering *BCL2L1* expression was more likely to correct the discordant predictions of **Cis1** (4 out of 5) than with **Cis2** (2 out of 4).

## DISCUSSION

Using gene expression signatures, we derived both GI_50_ threshold-dependent and -independent ML models which predict the chemotherapy responses for cisplatin, carboplatin and oxaliplatin, respectively. The cisplatin model **Cis1** (Supplementary Table S6A) most accurately predicted response in bladder cancer patients after 18 months, and **Car1** (Suppl. Table S6B) best predicted response in ovarian cancer patients after 4 years. **Oxa1** (Suppl. Table S6C) more accurately predicted disease-free patients than recurrent disease at the one year treatment threshold. The thresholds which best represented time-to-recurrence differed between the platin drugs in each cancer type. Cisplatin gene signatures had noticeably improved performance when smoking history was taken into account.

The three platin drugs produce distinctly different gene signature models. Initial gene sets exhibited some overlap between platin drugs (N=67 between any two platins), but very few of these were correlated by MFA of GI_50_ with multiple platin drugs (*ATP7B*, *BCL2* and *MSH2*). Signature genes common to multiple platin drugs whose expression was correlated with cisplatin GI_50_ values but not with carboplatin and/or oxaliplatin values include *BCL2L1, GSTP1, MAP3K1, MAPK3, MT1A,* and *MT2*. Similarly, genes correlating only to carboplatin GI_50_ included *AKT1, EGF, ERCC1, KRAS, LIG3, MTHFR, MTR, RAD50, TP53*, while genes correlating to only oxaliplatin GI_50_ included *ATM, BCL2, CLCN6, ERCC2, ERCC6,* and *UGT1A1*. Despite the close similarity between cisplatin and carboplatin GI_50_ response (see Figure 2), only one gene (*ATP7B*) was related by MFA to GI_50_ levels of both drugs. *BCL2* and *MSH2* correlated with both cisplatin and oxaliplatin GI_50_ (*BCL2* did not correlate with carboplatin GI_50_). The increase in misclassification caused by the elimination of *MSH2* from any SVM model in which it was present was significant; for example, misclassification of **Cis14** and **Oxa21** (Table 1) were increased by 28.2% and 19.1%, respectively (Suppl. Tables S2A and S2C). These differences may reflect the spectrum of activity, sensitivity, and toxicity of these signature genes^21–23,34,35^.

Previous validation of patient data for other drugs validated with other datasets^6,24^ using biochemically inspired machine learning have had better performance than those reported here. We investigated the possibility that disease and molecular heterogeneity in platin-treated patients may have affected the accuracy of our results. Model predictions were reevaluated after stratifying clinical features such as time-to-disease recurrence, cancer stage, and metastatic lymph node count. Breast cancer patients with advanced disease (stage III and IV) were analyzed separately from those with earlier stage diagnoses (stage I and II). Cisplatin model **Cis1** performed best on stage IV patients (overall accuracy 72.4% at a 2 year recurrence threshold), while **Oxa1** similarly performed best in predicting late stage cancers (74.5% accurate for stage III and 71.4% accurate for stage IV at a 2 year recurrence threshold). **Cis5** was also more accurate for later stage cancer patients (72.4% overall accuracy at 18 months). The accuracies of models were similar across all stages (e.g. **Car1** ranged from 58-74%). Cisplatin-treated, TCGA bladder cancer patients and oxaliplatin-treated TCGA colorectal cancer patients were also stratified by Lymph Node status (N0, N1, and N2 [bladder cancer patient data set comprised of only two N3 patients, which were included with the analysis of N2 patients; N3 was not represented in colorectal cancer]). In TCGA bladder cancer patients, **Cis1** exhibited ~60% accuracy across all categories; however it performed better in sensitive N0 and N1 patients relative to N2. **Cis2** was less accurate for N2 patients than for N0 and N1. Sensitive N2 patients were more likely to be misclassified (<40%) than relapsed N2 patients. In TCGA colorectal cancer patients, **Oxa1** was 88% accurate in N2 patients (95% accurate for sensitive N2 patients [n=19], and 67% accurate for relapsed N2 patients [n=6]). Oxaliplatin models were less accurate for N1 patients compared to N0 and N2. Thus, heterogeneity in disease stage as well as metastatic phenotypes adversely confounds the overall accuracies of our predictions.

Gene signature models derived from cell lines and tested on patients differ in their outcome measures. The exact GI_50_ cell line threshold that is most predictive of patient outcome is not known, and different groups use different methods to discretize GI_50_ values^36,37^. Therefore, we developed ML models for platin drugs which predict drug response without relying on arbitrary GI_50_ thresholds. For cisplatin, SVM ensemble averaging generated on different resistance thresholds shows a small increase in accuracy over most models, better representing the sensitive, disease-free class (59% accuracy). Interestingly, ensemble averaging of only the models built using a moderate GI_50_ thresholds yielded results which better represented the resistance class. This result closer matches the accuracy of **Cis1**, and may be due to **Cis1** having a greater overall impact on the ensemble prediction. When limiting ensemble averaging to only those models with the highest area under the curve (AUC) at each resistance threshold, differences in predictions were negligible. Ensemble ML can potentially avoid problems with poor performance and overfitting by combining models that individually perform slightly better than chance^38^.

It is difficult to reconcile gene signatures without features known to be related to chemoresistance with tumor biology. Our thresholding approach may reveal potentially important genes and pathways associated with platin resistance. It would be preferable to explore pathways related to signature genes to improve accuracy, identify potential targets for further study of chemoresistance, and expand the model parameters to take into account alternate states besides those captured in the original signature^39^. Signatures for resistance may be useful for developing targeted intervention to re-sensitize tumours. For example, the mismatch repair (MMR) gene *MSH2* is commonly present in gene signatures at high resistance levels for oxaliplatin, which is of interest, as MMR deficiency has been shown to be predictive for oxaliplatin resistance^35^. Indeed, *MLH1*, *MSH2* and *MSH6*-deficient cells are more susceptible to oxaliplatin, despite MMR-deficiency being associated with cisplatin resistance^34^. The autoimmune disease-associated gene *SIAE*, which has been previously shown to have a strong negative correlation to oxaliplatin response in advanced CRC patients^40^, was selected in the majority of thresholded oxaliplatin models (Supplementary Table S2C). The gene *BCL2*, which was commonly selected for cisplatin (Figure 3A), was rarely selected for oxaliplatin (Figure 3C). At the highest levels of resistance to cisplatin, models were enriched for genes belonging to DNA repair, anti-oxidative response, apoptotic pathways and drug transporters (Figure 3A). These gene pathways are known to be involved in cisplatin resistance^41,42^ and these specific genes may be explored in subsequent work to identify the contribution to chemotherapy response in a biochemical context.

Log-loss evaluates the accuracy of a classifier by penalizing erroneous classifications, and is relevant in cases where data is imbalanced and/or have an unequally distributed error cost. We assessed whether ML models based on log-loss minimization could improve model accuracy in patient data (Supplementary Table S4) and compared these to models generated by minimizing cell line misclassification. When models generated by both methods were highly similar (generated at the same GI_50_ threshold, consist of a similar number of genes and consist of ≥ 80% shared genes), prediction accuracy of TCGA cancer patient outcomes were nearly indistinguishable, as accuracy can vary over different relapse thresholds. Where significant differences in predictions were seen, the misclassification-based models were more accurate overall (**Cis1**, **Cis17** and the “12-Resistant” carboplatin model were +8.3%, +5.6% and +3.9% more accurate compared to the log-loss model, respectively). Oxaliplatin models were dissimilar across all GI_50_ thresholds, as the log-loss minimized ML models often contain increased numbers of genes compared to the misclassification-based models. Many of these larger models were less accurate in patients compared to models which minimized misclassification rates consistent that this evaluation and model selection method is more prone to overfitting. This pattern was also noted for models generated at extreme GI_50_ thresholds for all three platin drugs in which response was, by definition, somewhat imbalanced.

It may be feasible to predict responses to combination chemotherapy with the models described here. Not included in the present analysis were signatures for methotrexate, vinblastine, and doxorubicin, which comprise the MVAC cocktail used to treat bladder cancer. This was due primarily to a lack of patients treated with this drug combination in the TCGA bladder dataset (N=11). Individual signatures for several of these drugs have been derived and analyzed using the patient data from METABRIC (Molecular Taxonomy of Breast Cancer International Consortium)^24^. A reasonable approach to predicting combination chemotherapy would first determine the probability of sensitivity or resistance to individual drugs, accounting for the misclassification rate by each (defined as d_1_, …, d_k_). The ML classifiers output these probabilities, analogous to their misclassification rates in a set of patients treated identically. If the model predicts that the patient is sensitive to drug d_1_ with 90% probability, and sensitive to drug d_2_ with 5% probability, the probability of sensitivity to the combination is 1 - (1 - 0.9)*(1 - 0.05) = 90.5%, and the probability of resistance is 9.5%. The correlated responses could be estimated for drug pairs, d_1_ and d_2_, and then adjusted for the combined probability of the pair to d_12_, based on the features that are shared by the signatures of both drugs. The probability of sensitivity would then be given by 1 - (1 - d_12_)*(1 - d_3_)*…*(1 - d_k_).

The predictive accuracy for the same model could differentiate highly between the two datasets. **Cis3** (Supplemental Table S6A) had a high predictive accuracy and AUC for TCGA bladder cancer patients (AUC=0.64). However, the AUC was lower when applied to the Als *et al.*^29^ dataset (AUC=0.18). Patient metadata in the latter study only indicated patient survival times, while we base the expected TCGA patient outcome on time to disease recurrence. As the basis of our expected outcome differs between datasets, these differences may be acting as a confounding factor to determine accuracy of gene signatures. The datasets also differ in how expression was measured (microarray vs. RNA-seq). The relevance of models based on training and testing data from different platforms can affect the accuracy of validation, which might not be improved by data normalization. In this study, datasets were subjected to z-score normalization. In subsequent studies, other techniques to correct for some of these effects have been described and could be applied^43^.

In summary, we describe GI_50_-or IC_50_-threshold-independent ML models to predict chemotherapy response to platin agents in cancer patients. Ensemble machine learning produced combined signatures that were more accurate than most individual models generated with different thresholds. Genes associated cisplatin response included those which exacerbate resistance in patients with a history of smoking. The methodology described here should be adaptable to other drugs and cancer types. With a range of models for multiple drugs, it may be possible to improve the efficacy of treatment by tailoring treatment to a patient’s specific tumour biology, and reduce treatment duration by limiting the number of different therapeutic regimens prescribed before achieving a successful response^44^.

## MATERIALS AND METHODS

### Data and preprocessing

Microarray GE and data were from breast cancer cell lines were used to train ML-based gene signatures of drug response based on respective growth or target inhibition data (GI_50_ or IC_50_). Cell lines were treated with either cisplatin (N=39), carboplatin (N=46), or oxaliplatin (N=47)^13^. Bladder cancer cell line GE and IC_50_ measurements for cisplatin were obtained from cancerRxgene (N=17). However, all testing was performed on breast cancer cell line data as the number of bladder cancer cell lines was insufficient to produce accurate signatures. RNA-seq GE and survival measurements were downloaded from TCGA for bladder urothelial carcinoma (N=72 patients treated with cisplatin)^26^, ovarian epithelial tumor (N=410 treated with carboplatin)^27^ and colorectal adenocarcinoma (N=99 treated with oxaliplatin)^28^. GE of cisplatin-treated patients of cell carcinoma of the urothelium (N=30)^29^ and for oxaliplatin-treated CRC patients (N=83)^30^ were obtained from the Gene Expression Omnibus. Clinical metadata and GE for TCGA patients were obtained from Genomic Data Commons (https://gdc.cancer.gov/), while methylation HM450 (Illumina) data for these patients was downloaded from cBioPortal^45^.

Initial gene sets for developing signatures for each drug were identified from previously published literature (see Supplemental References) and databases, such as PharmGKB and DrugBank^46,47^. The final gene sets were chosen using MFA to analyze interactions between GE, CN, and GI_50_ data for the drug of interest^48^. Genes whose GE and/or CN showed a direct or inverse correlation with GI_50_ were selected for SVM training. As the number of genes related to GI_50_ for oxaliplatin exceeded the number of cell lines available for training, we limited the input to the ML model oxaliplatin to those genes whose GE were related to GI_50_. Similarly, the number correlating genes in cisplatin treated cells exceeded the number of cell lines. For cisplatin, genes whose expression correlated with GI_50_ were eliminated if they showed no or little expression in TCGA bladder cancer patients (i.e. RNA-seq counts by Expectation Maximization [RSEM] were < 5.0 for majority of individuals). This reduces the overall number of genes for SVM analysis, and thus helps to avoid a data to size sample imbalance. For cisplatin, MFA was repeated using IC_50_ values for 17 bladder cancer cell lines; however, the available CN data generally showed a lack of variation in the cell lines for these genes. Instead, the available IC_50_ values for three other cancer drugs (doxorubicin, methotrexate and vinblastine) were compared with the IC_50_ of cisplatin by MFA.

Applying an SVM model directly to patient data without a normalization approach is imprecise when training and testing data are not obtained using similar methodology (i.e. different microarray platforms). To compare the cell line GE microarray data and the patient RNA-seq GE datasets, expression values were normalized by conversion to z-scores using MATLAB^49^. Although Log2 intensity values from microarray data were not available for TCGA samples, RNA-seq based GE and log_2_ intensities from microarray data are highly correlated^50^.

### Machine Learning

SVMs were trained with breast cancer cell line GE datasets^13^ with the Statistics Toolbox in MATLAB^49^ similar to Dorman et al (2016)^6^. Rather than a linear kernel, we used a Gaussian kernel function (fitcsvm), and then tested with leave-one-out cross-validation (using the options ‘*crossval’* and ‘*leaveout’*). A greedy backwards feature selection algorithm was used to improve classification accuracy^51^. BFS leaves out individual genes from the initial MFA-qualified gene set, then trains a cross validated Gaussian kernel SVM on the training samples, removing the gene with the highest misclassification rate. The procedure is repeated until all genes have been evaluated. The gene subset with the lowest misclassification rate^6^ or log-loss statistic^25^ based on cross-validation is selected as the model for subsequent testing with patient GE and clinical data. K-fold cross validation of the misclassification-based models was performed using MATLAB software described in Zhao et al. (2018)^25^.

SVMs minimized according to the log-loss classification function were also generated with both software described in Zhao *et al.* (2018; uses multiclass compatible ‘fitcecoc’ function)^25^, and with a modified version of the software described above (using ‘fitSVMPosterior’ to compute posterior probabilities). Computed probabilities differ between ‘fitSVMPosterior’ and ‘fitcecoc’ (range: 0.02-0.04), thus the resultant models will differ between the two programs. When given unbalanced data (e.g. lower resistance thresholds), ‘fitSVMPosterior’ will warn that some classes are not represented, and thus those folds will not predict the labels for those missing classes. The log-loss models described in this manuscript were generated with the multiclass compatible ‘fitcecoc’ function software^25^.

#### Derivation of gene signatures for different drug resistance thresholds

We have previously set a conventional GI_50_ threshold distinguishing sensitivity from resistance at the *median* of the range of drug concentrations that inhibited cell growth by 50%^6^. We hypothesized that different gene signatures could be derived for different levels of drug resistance by varying this threshold. ML experiments for classifying resistance or sensitivity at GI_50_ values generated a series of optimized Gaussian SVM models whose performance were assessed with patient expression data for each signature. A heat map which illustrates the frequencies of genes appearing in these models was created with the R language *hist2d* function.

A composite gene signature was created by ensemble averaging of all models generated at each resistance threshold. Ensemble averaging combines signatures through averaging the weighted accuracy of a set of related models^38^. The decision function for the ensemble classifier is the mean of the decision function scores of the component classifiers, weighted by the AUC.

#### Significance of cell line-derived models

The potential for models to overfit data during training and/or feature selection was first assessed by permutation analysis with randomized cell line labels and with random sets of genes, as described previously^6^. Using the median cisplatin GI_50_ as the resistance threshold, 10,000 models based on random gene selection (15 genes) had higher rates of misclassification than the best median SVM models (2 signatures with 7.7% misclassification). Cisplatin, carboplatin and oxaliplatin GE data for random cell line label combinations (n=10,000) generated only 8, 1 and 1 signatures, respectively, with lower error rates than the best biochemically-inspired signatures. When minimizing for log-loss (rather than misclassification), random gene analysis (10,000 iterations; median cisplatin GI_50_ threshold) resulted only in models with a higher log-loss than the model generated with the initial cisplatin gene set. Log-loss based random label analysis (n=2000 combinations) resulted in 3.4% of random label models resulted in a lower log-loss than the cisplatin model at the same GI_50_ threshold (5.27).This was not entirely surprising, since this result depends on the GI_50_ threshold used for labeling. The differences between GI_50_ values for cell lines close to the median GI_50_ used in this analysis are almost negligible (e.g. 5.11 vs 5.12) and likely within the measurement error for these values.

Cell-line based model accuracies in predicting outcomes of platin-treated TCGA bladder cancer patients were compared with results from participants who did not receive these treatments (using an 18 months post-treatment threshold). In non-platin treated patients, 36.5% of those who were disease-free were predicted accurately with the **Cis1** signature (N=178; 22% less accurate than platin treated patients), and 62.9% accurate for those with recurrent disease (N=70; 8.1% less accurate). **Cis2** was 43.8% accurate for disease-free non-platin treated patients (N=178; 12.3% lower accuracy), and 60.0% of those who relapsed (N=70; 2.9% less accurate). Gene expression changes in patients treated with platin drugs are better modeled by cancer cell-line based predictors than in patients receiving other treatments.

## ACKNOWLEDGEMENTS

Katherina Baranova contributed to the cisplatin gene signatures and Dimo Angelov developed automated feature selection. We thank Murray Junop for commenting on the manuscript. Compute Canada and Shared Hierarchical Academic Research Computing Network (SHARCNET) provided high performance computing and storage facilities.

## AUTHOR CONTRIBUTIONS

PKR and DL designed the methodology. EJM and JZ performed analyses. EJM and PKR wrote the manuscript.

---

**Supplementary Figure 1.**
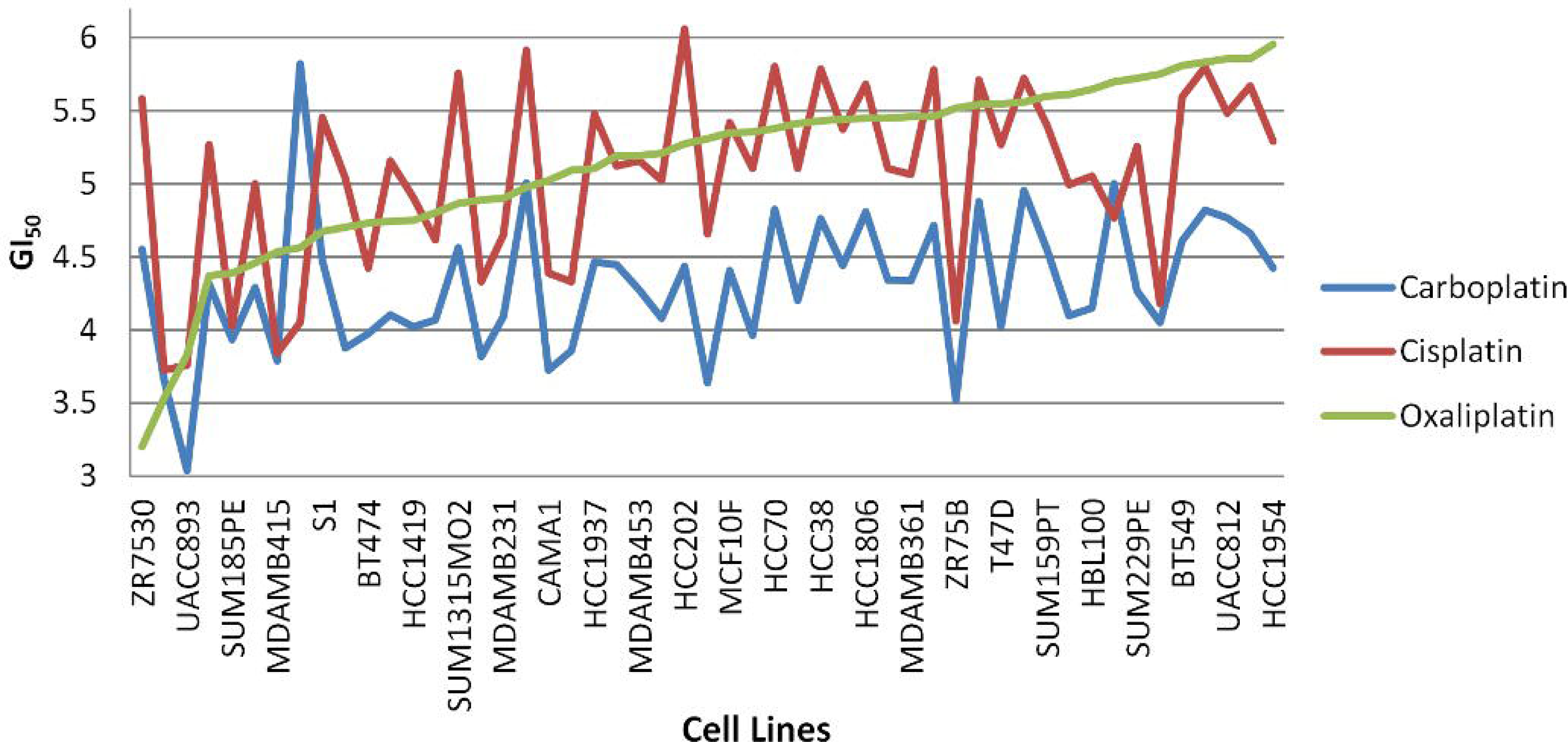
The variation in gene composition of log-loss based SVMs at different GI_50_ thresholds for A) cisplatin, B) carboplatin, and C) oxaliplatin. Each box represents the density of genes appearing in optimized Gaussian log-loss SVM models in those functional categories, with darker grey indicating frequent genes in indicated GI_50_ threshold intervals, while lighter grey indicates less commonly selected genes.

**Supplementary Figure 2.**
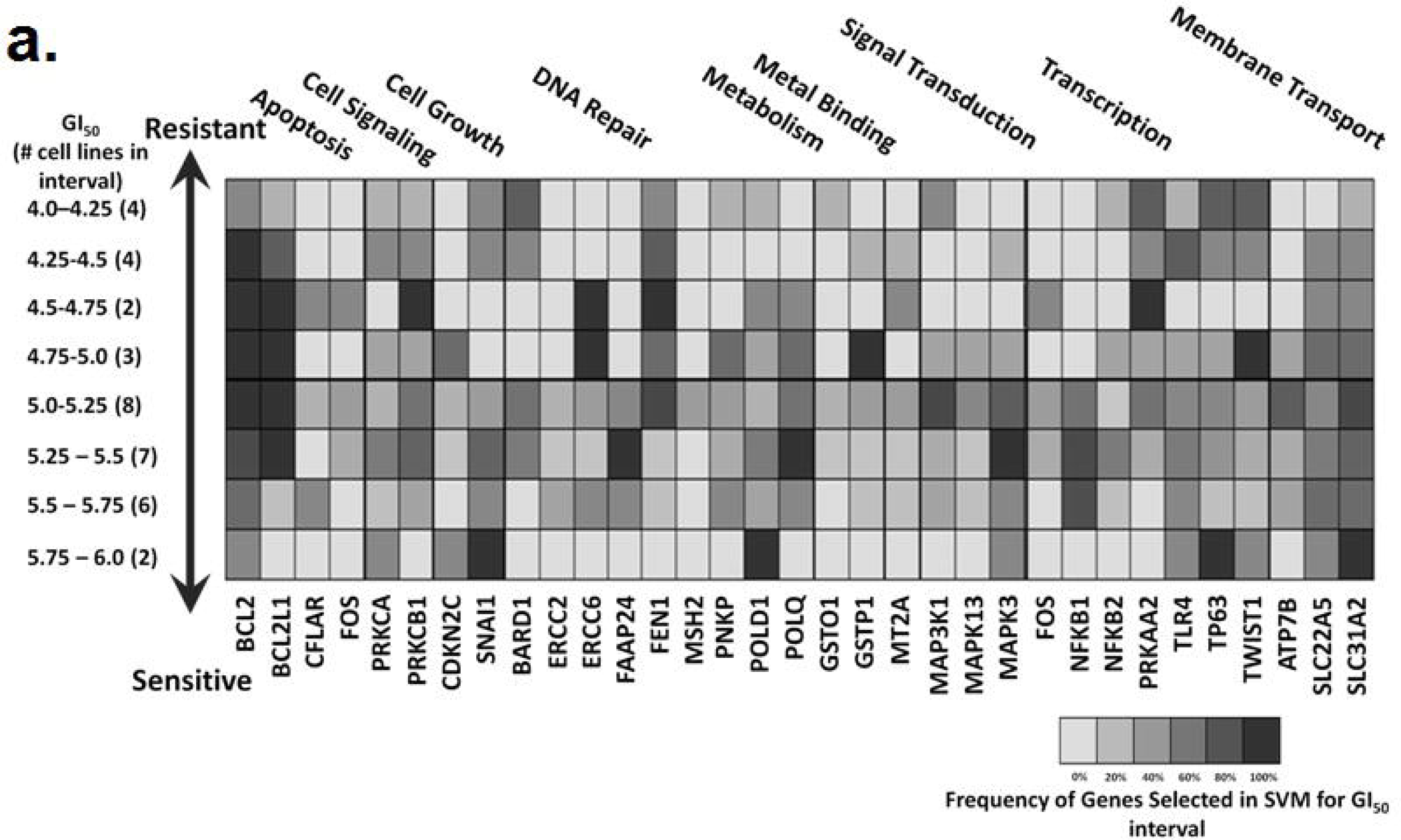
Classification accuracy of models on TCGA bladder cancer patients treated with cisplatin and/or carboplatin as the resistance threshold is varied. Recurrence and disease-free survival are used as a binary measure to assess performance. The x-axis indicates movement of the resistance threshold, with more cell lines labeled sensitive on the left and more labeled resistant on the right. Maximal AUC is indicated by the downward arrows.

**Supplementary Figure 3.**
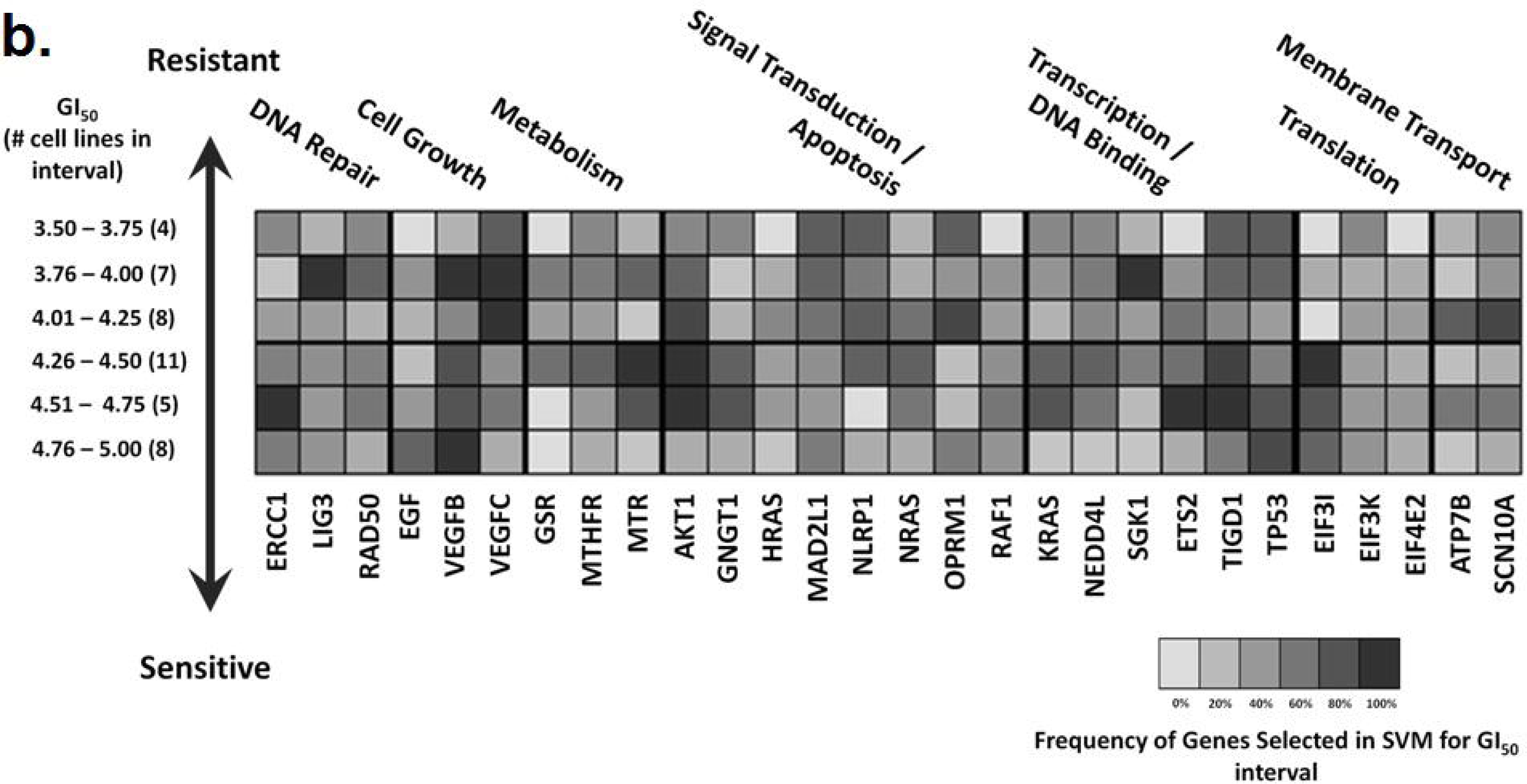
Classification accuracy of SVM models for cisplatin, at a range of response thresholds, were assessed using gene expression data for cisplatin-treated bladder cancer patients from Als *et al.* ^29^. Patients with a ≥ 5 year survival post-treatment were labeled sensitive. Red arrows indicate the SVM models with the highest positive predictive value (PPV) in the accuracy of classification of patient outcome.

**Supplementary Figure 4.**
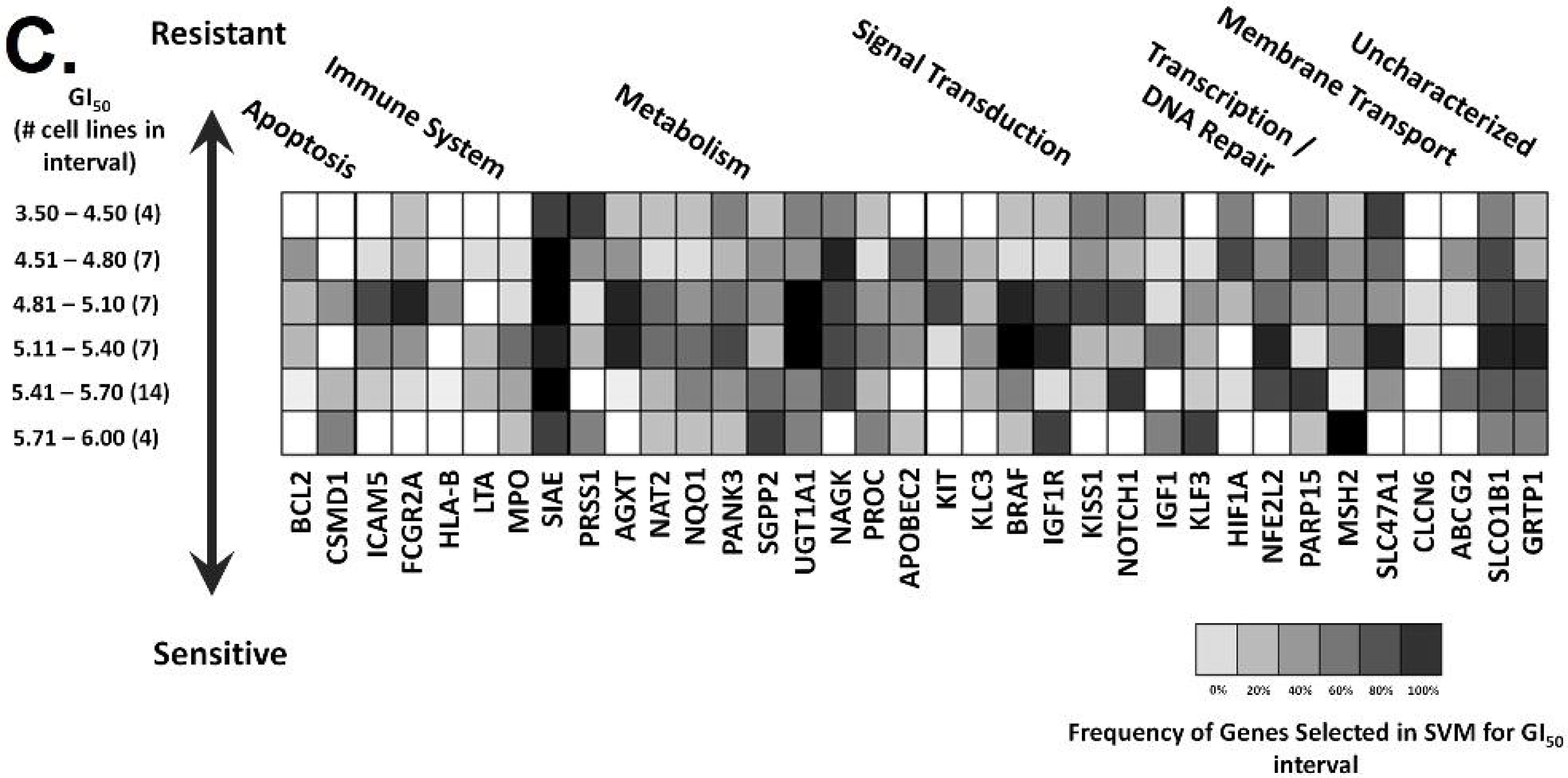
Hyperplane distance calculated by all thresholded SVMs for recurrent (<6 months) TCGA patients. Each diagram represents the predictions of all SVMs for all patients who had recurrence less than 6 months after treatment (N=10). Each point represents an SVM, where the x-axis represents the number of cell lines set to resistant (in order of lowest to highest GI_50_), and the y-axis represents the calculated hyperplane distance. A negative hyperplane distance would represent a prediction of resistance to cisplatin. Despite this, some patients show a strong preference towards predictions of sensitivity (i.e. TCGA-XF-A9SU).

**Supplementary Figure 5.** Hyperplane distance calculated by all thresholded SVMs for sensitive TCGA patients. Each diagram represents the predictions of all SVMs for all patients who had recurrence > 4 years after treatment (top; N=3), or patients who showed no recurrence after 6 years (bottom; N=6). Each point represents an SVM, where the x-axis represents the number of cell lines set to resistant (in order of lowest to highest GI_50_), and the y-axis represents the calculated hyperplane distance. A positive hyperplane distance would represent a prediction of sensitivity to cisplatin.

## REFERENCES

1. Cardoso, F. et al. Locally recurrent or metastatic breast cancer: ESMO Clinical Practice Guidelines for diagnosis, treatment and follow-up. Ann. Oncol. 23, vii11–vii19 (2012).

2. Oostendorp, L. J., Stalmeier, P. F., Donders, A. R. T., van der Graaf, W. T. & Ottevanger, P. B. Efficacy and safety of palliative chemotherapy for patients with advanced breast cancer pretreated with anthracyclines and taxanes: a systematic review. Lancet Oncol. 12, 1053–1061 (2011).

3. Alfarouk, K. O. et al. Resistance to cancer chemotherapy: failure in drug response from ADME to P-gp. Cancer Cell Int. 15, 71 (2015).

4. Gąsowska-Bodnar, A. et al. Survivin Expression as a Prognostic Factor in Patients With Epithelial Ovarian Cancer or Primary Peritoneal Cancer Treated With Neoadjuvant Chemotherapy: Int. J. Gynecol. Cancer 24, 687–696 (2014).

5. Hatzis, C. et al. A genomic predictor of response and survival following taxane-anthracycline chemotherapy for invasive breast cancer. JAMA 305, 1873–1881 (2011).

6. Dorman, S. N. et al. Genomic signatures for paclitaxel and gemcitabine resistance in breast cancer derived by machine learning. Mol. Oncol. 10, 85–100 (2016).

7. Zhang, S. et al. Organic Cation Transporters Are Determinants of Oxaliplatin Cytotoxicity. Cancer Res. 66, 8847–8857 (2006).

8. Poisson, L. M. et al. A metabolomic approach to identifying platinum resistance in ovarian cancer. J. Ovarian Res. 8, (2015).

9. Cadoná, F. C. et al. Guaraná a Caffeine-Rich Food Increases Oxaliplatin Sensitivity of Colorectal HT-29 Cells by Apoptosis Pathway Modulation. Anticancer Agents Med. Chem. 16, 1055–1065 (2016).

10. Kasparkova, J., Vojtiskova, M., Natile, G. & Brabec, V. Unique Properties of DNA Interstrand Cross-Links of Antitumor Oxaliplatin and the Effect of Chirality of the Carrier Ligand. Chem. –Eur. J. 14, 1330–1341 (2008).

11. Woynarowski, J. M. et al. Oxaliplatin-Induced Damage of Cellular DNA. Mol. Pharmacol. 58, 920–927 (2000).

12. Tashiro, T., Kawada, Y., Sakurai, Y. & Kidani, Y. Antitumor activity of a new platinum complex, oxalato (trans-l-1,2-diaminocyclohexane)platinum (II): new experimental data. Biomed. Pharmacother. 43, 251–260 (1989).

13. Daemen, A. et al. Modeling precision treatment of breast cancer. Genome Biol. 14, R110 (2013).

14. Yuan, Y. et al. Identification of the biomarkers for the prediction of efficacy in first-line chemotherapy of metastatic colorectal cancer patients using SELDI-TOF-MS and artificial neural networks. Hepatogastroenterology. 59, 2461–2465 (2012).

15. L’Espérance, S., Bachvarova, M., Tetu, B., Mes-Masson, A.-M. & Bachvarov, D. Global gene expression analysis of early response to chemotherapy treatment in ovarian cancer spheroids. BMC Genomics 9, 99 (2008).

16. Nickerson, M. L. et al. Molecular analysis of urothelial cancer cell lines for modeling tumor biology and drug response. Oncogene (2016).

17. Yuryev, A. Gene expression profiling for targeted cancer treatment. Expert Opin. Drug Discov. 10, 91–99 (2015).

18. Sataloff, D. M. et al. Pathologic response to induction chemotherapy in locally advanced carcinoma of the breast: a determinant of outcome. J. Am. Coll. Surg. 180, 297–306 (1995).

19. Ogston, K. N. et al. A new histological grading system to assess response of breast cancers to primary chemotherapy: prognostic significance and survival. Breast Edinb. Scotl. 12, 320–327 (2003).

20. Earl, J. et al. The UBC-40 Urothelial Bladder Cancer cell line index: a genomic resource for functional studies. BMC Genomics 16, 403 (2015).

21. Rixe, O. et al. Oxaliplatin, tetraplatin, cisplatin, and carboplatin: Spectrum of activity in drug-resistant cell lines and in the cell lines of the national cancer institute’s anticancer drug screen panel. Biochem. Pharmacol. 52, 1855–1865 (1996).

22. Mehmood, R. K. Review of Cisplatin and oxaliplatin in current immunogenic and monoclonal antibody treatments. Oncol. Rev. 8, 256 (2014).

23. Kweekel, D. M., Gelderblom, H. & Guchelaar, H.-J. Pharmacology of oxaliplatin and the use of pharmacogenomics to individualize therapy. Cancer Treat. Rev. 31, 90–105 (2005).

24. Mucaki, E. J. et al. Predicting Outcomes of Hormone and Chemotherapy in the Molecular Taxonomy of Breast Cancer International Consortium (METABRIC) Study by Biochemically-inspired Machine Learning. F1000Research 5, 2124 (2017).

25. Zhao, J. Z. L., Mucaki, E. J. & Rogan, P. K. Predicting ionizing radiation exposure using biochemically-inspired genomic machine learning. F1000Research 7, 233 (2018).

26. Robertson, A. G. et al. Comprehensive Molecular Characterization of Muscle-Invasive Bladder Cancer. Cell 171, 540–556.e25 (2017).

27. Cancer Genome Atlas Research Network. Integrated genomic analyses of ovarian carcinoma. Nature 474, 609–615 (2011).

28. Cancer Genome Atlas Network. Comprehensive molecular characterization of human colon and rectal cancer. Nature 487, 330–337 (2012).

29. Als, A. B. et al. Emmprin and survivin predict response and survival following cisplatin-containing chemotherapy in patients with advanced bladder cancer. Clin. Cancer Res. Off. J. Am. Assoc. Cancer Res. 13, 4407–4414 (2007).

30. Tsuji, S. et al. Potential responders to FOLFOX therapy for colorectal cancer by Random Forests analysis. Br. J. Cancer 106, 126–132 (2012).

31. Tembe, V. et al. The BARD1 BRCT domain contributes to p53 binding, cytoplasmic and mitochondrial localization, and apoptotic function. Cell. Signal. 27, 1763–1771 (2015).

32. Freedman, N. D., Silverman, D. T., Hollenbeck, A. R., Schatzkin, A. & Abnet, C. C. Association between smoking and risk of bladder cancer among men and women. JAMA 306, 737–745 (2011).

33. Joehanes, R. et al. Epigenetic Signatures of Cigarette Smoking. Circ. Cardiovasc. Genet. (2016).

34. Raymond, E., Faivre, S., Chaney, S., Woynarowski, J. & Cvitkovic, E. Cellular and Molecular Pharmacology of Oxaliplatin. Mol. Cancer Ther. 1, 227–235 (2002).

35. Alex, A. K. et al. Response to Chemotherapy and Prognosis in Metastatic Colorectal Cancer With DNA Deficient Mismatch Repair. Clin. Colorectal Cancer (2016).

36. Sos, M. L. et al. Predicting drug susceptibility of non-small cell lung cancers based on genetic lesions. J. Clin. Invest. 119, 1727–1740 (2009).

37. Laderas, T. G., Heiser, L. M. & Sönmez, K. A Network-Based Model of Oncogenic Collaboration for Prediction of Drug Sensitivity. Front. Genet. 6, 341 (2015).

38. Clemen, R. T. Combining forecasts: A review and annotated bibliography. Int. J. Forecast. 5, 559–583 (1989).

39. Airley, R. Cancer chemotherapy. (Wiley-Blackwell, 2009).

40. Li, X.-X. et al. RNA-seq identifies determinants of oxaliplatin sensitivity in colorectal cancer cell lines. Int. J. Clin. Exp. Pathol. 7, 3763–3770 (2014).

41. Borst, P., Rottenberg, S. & Jonkers, J. How do real tumors become resistant to cisplatin? Cell Cycle Georget. Tex 7, 1353–1359 (2008).

42. Wernyj, R. & Morin, P. Molecular mechanisms of platinum resistance: still searching for the Achilles? heel. Drug Resist. Updat. 7, 227–232 (2004).

43. Johnson, W. E., Li, C. & Rabinovic, A. Adjusting batch effects in microarray expression data using empirical Bayes methods. Biostat. Oxf. Engl. 8, 118–127 (2007).

44. Akamatsu, N., Nakajima, H., Ono, M. & Miura, Y. Increase in acetyl CoA synthetase activity after phenobarbital treatment. Biochem. Pharmacol. 24, 1725–1727 (1975).

45. Gao, J. et al. Integrative analysis of complex cancer genomics and clinical profiles using the cBioPortal. Sci. Signal. 6, pl1 (2013).

46. Whirl-Carrillo, M. et al. Pharmacogenomics Knowledge for Personalized Medicine. Clin. Pharmacol. Ther. 92, 414–417 (2012).

47. Law, V. et al. DrugBank 4.0: shedding new light on drug metabolism. Nucleic Acids Res. 42, D1091–D1097 (2014).

48. Abdi, H. & Williams, L. J. Principal component analysis. Wiley Interdiscip. Rev. Comput. Stat. 2, 433–459 (2010).

49. MATLAB and Statistics Toolbox Release 2012b, The MathWorks, Inc., Natick, Massachusetts, United States.

50. Marioni, J. C., Mason, C. E., Mane, S. M., Stephens, M. & Gilad, Y. RNA-seq: an assessment of technical reproducibility and comparison with gene expression arrays. Genome Res. 18, 1509–1517 (2008).

51. Bermingham, M. L. et al. Application of high-dimensional feature selection: evaluation for genomic prediction in man. Sci. Rep. 5, 10312 (2015).

